# Skeletal descriptions of shape provide unique perceptual information for object recognition

**DOI:** 10.1101/518795

**Authors:** Vladislav Ayzenberg, Stella F. Lourenco

**Affiliations:** Department of Psychology, Emory University

## Abstract

With seemingly little effort, humans can both identify an object across large changes in orientation and extend category membership to novel exemplars. Although researchers argue that object shape is crucial in these cases, there are open questions as to how shape is represented for object recognition. Here we tested whether the human visual system incorporates a three-dimensional skeletal descriptor of shape to determine an object’s identity. Skeletal models not only provide a compact description of an object’s global shape structure, but also provide a quantitative metric by which to compare the visual similarity between shapes. Our results showed that a model of skeletal similarity explained the greatest amount of variance in participants’ object dissimilarity judgments when compared with other computational models of visual similarity (Experiment 1). Moreover, parametric changes to an object’s skeleton led to proportional changes in perceived similarity, even when controlling for another model of structure (Experiment 2). Importantly, participants preferentially categorized objects by their skeletons across changes to local shape contours and non-accidental properties (Experiment 3). Our findings highlight the importance of skeletal structure in vision, not only as a shape descriptor, but also as a diagnostic cue of object identity.

## Introduction

The same object produces vastly different shapes on the retina across changes in orientation, and objects of the same category have vastly different shape contours across exemplars. Yet, with very little experience, humans (adults^1^, infants^2^) and nonhuman animals (monkeys^3^, chicks^4^, rats^5^) recognize objects rapidly and with ease across such variations. Research in the vision sciences suggests that shape is crucial for object recognition^6–8^. Humans readily use shape to recognize objects in the absence of other visual information (e.g., texture and shading) ^7,9^ and both adults and children preferentially categorize novel objects by their shape across conflicting color and texture cues^10,11^. Moreover, human representations of shape are robust to changes in view^1,12^, contour perturbation^13,14^, and deformations from bending or stretching (e.g., hand poses)^15–17^, suggesting a reliance on global shape properties over local contour information^18,19^. However, few studies have offered a formalized account of how humans represent and compare global shape properties to recognize objects^6^. In the current study, we tested whether humans incorporate a computational model of global shape structure based on the medial axis, or ‘shape skeleton’, to determine an object’s identity.

Shape skeletons are a class of geometric models, based on the medial axis of the shape. These models describe shape via the set of symmetry axes that lie equidistant between two or more points along the boundary^20,21^ (see Figure 1). For most shapes, the axes are organized hierarchically, such that there may be a series of parent axes that describe the shape’s coarse global geometry, as well as smaller ‘off-shoot’ axes that describe individual component parts. More specifically, skeletons describe an object’s shape structure by specifying the spatial configuration of contours and component parts. Modern skeletal algorithms (i.e., pruned medial axis models^22,23^) are particularly good descriptors of an object’s global shape because their structure remains relatively stable across contour variations typical of natural contexts (e.g., perturbations, bending)^24–26^. Importantly, research in computer vision has formalized many methods by which to compare skeletons (e.g., distance metrics^27^), thereby providing a quantitative metric by which to compare shapes. These methods have been used successfully to identify objects across viewpoints and exemplars^28^, with classification accuracy matching human performance for superordinate object categories^29^.

**Figure 1.**
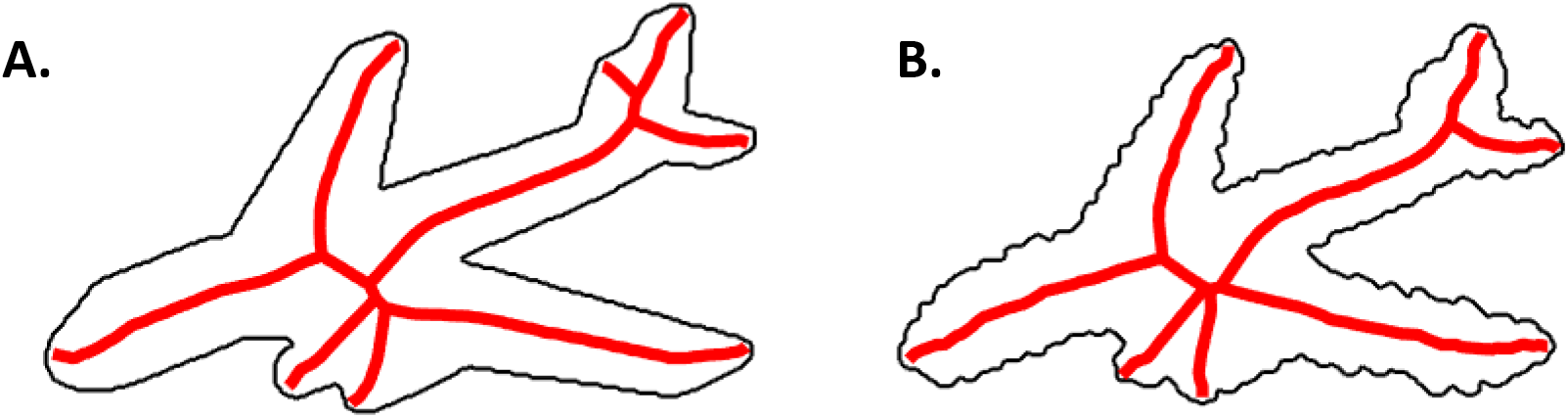
An illustration of the shape skeleton for a 2D airplane with (B) and without (A) perturbed contours. A strength of a skeletal model is that it can describe an object’s global shape structure across variations in contour. Skeletons computed using the ShapeToolbox^24^.

Accumulating evidence suggests that the skeletal structure of objects is extracted by the primate visual system during shape perception. In particular, behavioral studies have shown that human participants extract the skeletons of different 2D shapes^30–34^ and those skeletal structures remain relatively stable across border disruptions resulting from perturbations or illusory contours^35^. Increasingly, studies have shown that skeletal structures may be represented in three dimensions (3D) within an object-centered reference frame. Indeed, humans are better at discriminating 3D objects by skeletal differences than by differences in component parts (e.g., part orientation)^36^. Moreover, studies using neural recording (i.e., fMRI and electrophysiology) with humans and monkeys have found sensitivity to 3D object skeletons in high-level visual cortical areas (e.g., IT), including those known to support object recognition^37,38^. Skeletal sensitivity in these regions was decoded across changes in orientation and variations in local shape properties, suggesting a 3D object-centered representation that is robust to changes in viewpoint and component parts.

Despite the success of skeletal descriptions in computer vision systems^26^ and their biological plausibility in the primate visual system^37^, shape skeletons are rarely incorporated into models of object recognition. Instead, modern computational approaches to object recognition emphasize image statistics^39^ or hierarchical feature extraction operations such as those implemented by convolutional neural networks (CNNs)^40,41^. Yet, without explicitly invoking any skeletal description, these models match human performance on object recognition tasks^41^, and they are predictive of both human behavioral and neural responses^42–44^. Even models that do emphasize global shape properties do so by describing the local properties of components parts (e.g., geons) and coarse, categorically-defined, spatial relations^12,45^, not a skeletal structure. Given that these other models successfully approximate human object recognition, one might ask whether skeletal descriptions of shape are necessary for human object recognition at all. Thus, in the current study, we tested the degree to which skeletal descriptions of shape make unique, and possibly privileged, contributions to human object recognition in comparison to several other models of shape and object perception.

If the shape’s skeletal structure provides unique contributions to object recognition, then humans should perceive objects with similar skeletons as more similar to one another, even when controlling for other models. Moreover, if skeletal structures are a privileged source of information for object recognition, then humans should favor the shape skeleton over both non-shape based models of visual similarity, as well as other descriptors of shape. To this end, we assessed whether participants’ perceptual judgments of object similarity scaled with the skeletal similarity between novel 3D objects (Experiment 1), including objects whose coarse spatial relations could not be used for judging similarity (Experiment 2). We also tested how participants classified objects when the shape’s skeletal structure was placed in conflict with the object’s surface form, a manipulation that altered the shape’s contours and non-accidental properties (NAPs) without changing its skeleton (Experiment 3). In all cases, we examined the unique contributions of skeletal structures in object recognition by contrasting the shape skeleton with models of vision that do not explicitly incorporate a skeletal structure, but are nevertheless predictive of human object recognition. These models included those that describe visual similarity by their image statistics, namely, the Gabor-Jet (GBJ) model^46^ and GIST model^47^, as well as biologically plausible neural networks models, namely, the HMAX model^40^ and AlexNet, a CNN pre-trained to identify objects^41^. To anticipate our findings, a model of skeletal similarity was predictive of participants’ perceptual similarity and classification judgments even when accounting for these other models, suggesting that skeletal descriptions of shape play a crucial role in human object recognition, independent of other models of shape and object perception.

### Experiment 1 – Is perceived object similarity uniquely predicted by a model of skeletal similarity?

Here we tested one of the predictions outlined above: namely, as object skeletons become more similar, participants should judge the objects as being more alike. To test for a relation between human perceptual judgments and the shape skeleton, we generated a novel set of 3D objects that varied in their skeletal structures. Crucially, we compared the predictive power of skeletal descriptions to other models of visual similarity and tested the degree to which a model of skeletal similarity explained unique variance in human perceptual judgments.

#### Stimuli and experimental design

A total of 150 3D objects consisting of 30 skeletons were generated (see Figure 2A). All objects were comprised of three segments and were normalized for overall size (see Methods). Each object was rendered with five surface forms, serving to change the visible shape of the object on the retina without altering the underlying skeleton (see Figure 2B and Methods). Skeletal similarity between every object was calculated in 3D, object-centered, space as the mean Euclidean distance between each point on one skeleton and the closest point on the second skeleton following maximal alignment (see Methods). We chose to test a 3D skeletal description because of behavioral^48^ and neural^49^ evidence for 3D object-centered representations in the visual system, which include a sensitivity to 3D skeletal structures^36,37^.

**Figure 2.**
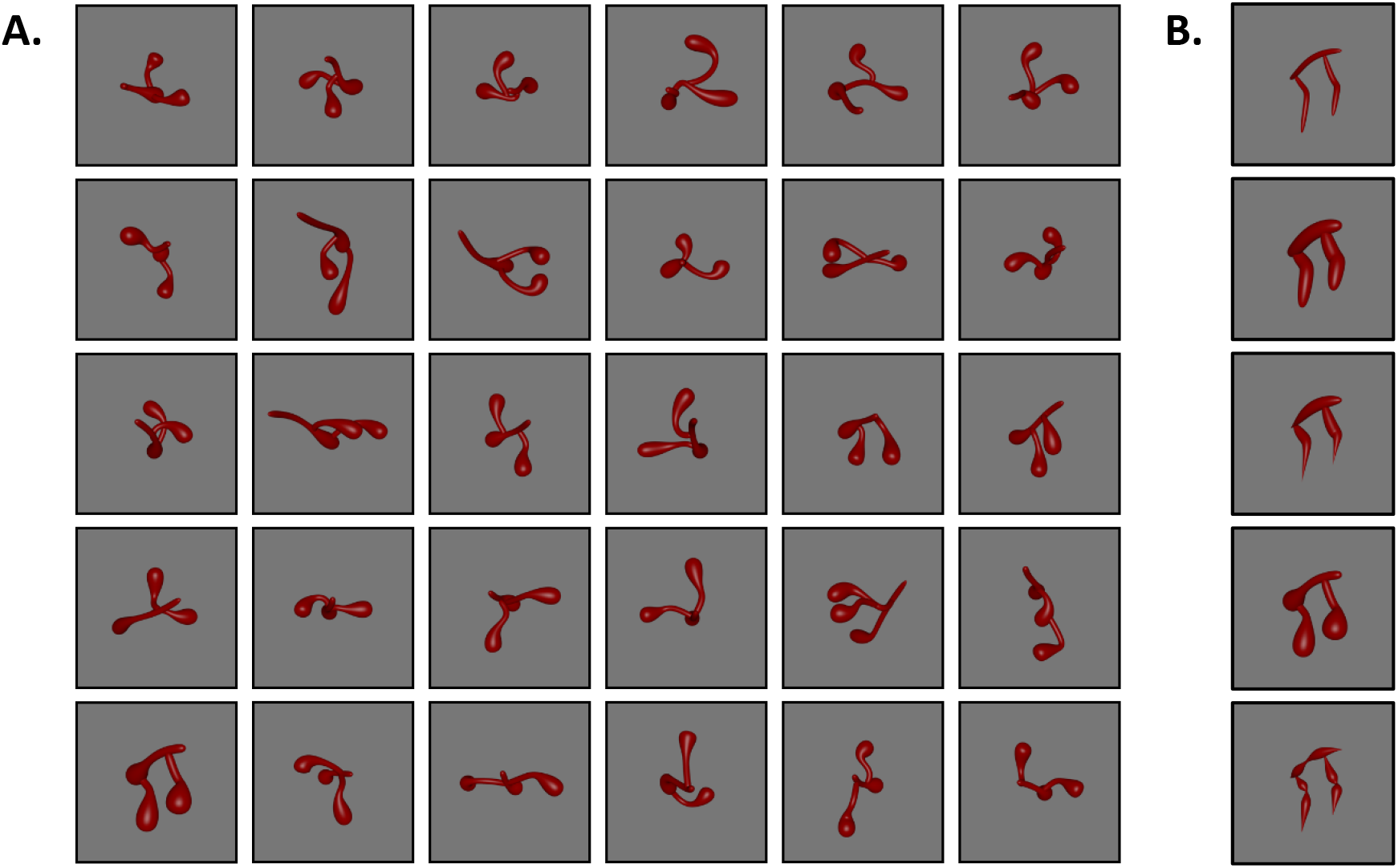
Stimuli used in Experiment 1. (A) Objects were procedurally generated to have different skeletal structures. (B) Each object was also rendered with five surface forms so as to vary in contour shape and non-accidental properties (NAPs) without disrupting the object’s skeleton. A cluster analysis revealed that the first and second surface form (from top to bottom) were comprised of the same NAPs (see Experiment 3 and Methods for more stimulus details). Subsets of these stimuli were used in Experiments 2 and 3 (see Methods).

Participants (*n* = 42) were administered a discrimination task in which they were shown images of two objects presented simultaneously in one of three depth orientations (−30°, 0°, +30°), with either the same or different skeletons. Participants were instructed to decide whether the two images showed the same or different object (independent of orientation). Participants were given unlimited time to respond but the instructions emphasized speed and accuracy.

We chose to use an untimed discrimination task where the objects were presented simultaneously in order to minimize task demands. However, we also confirmed that this task could be accomplished in a speeded context and found comparable performance to that reported below (see Supplemental Experiment 1).

#### Results and discussion

Participants discriminated the objects significantly above chance (0.50) *M*_*accuracy*_ = 0.80, *t*(41) = 17.64, *p* < .001, *d* = 2.72 (*M*_*RT*_ = 2129 ms). Thus, even though our stimulus set may be considered one class of object, and potentially difficult to discriminate, the objects differed sufficiently to support accurate discrimination (see also Supplemental Experiment 1 for comparable performance in a speeded version of the task).

To analyze whether a skeletal model was predictive of human object judgments, we converted participants’ binary discrimination judgments for each object pair into a continuous dissimilarity score. Dissimilarity scores for each object pair were computed by taking the mean discrimination accuracy for each pair across all participants. Human judgments were compared to each model by regressing human dissimilarity scores on model dissimilarity scores (see Methods).

Skeletal similarity was a significant predictor of participants’ judgments, *r* = 0.30, *p* < 0.001, explaining 9% of the variance (significance determined via permutation test; see Figure 3). That is, as the similarity between skeletal structures increased, participants were more likely to judge the objects as the same. However, one might ask whether another model of vision, which does not incorporate skeletal information, would also correlate with human judgments. To answer this question, we compared participants’ judgments to GBJ, GIST, HMAX, and AlexNet models. When compared independently, these models were all predictive of participants’ judgments (*rs* = 0.25 – 0.32, *r*^2^ = 6% – 11%; see Figure 3), with no significant differences between models (overlapping confidence intervals). For context, a noise ceiling representing a hypothetical true model (calculated by repeatedly splitting participants’ data into two sets and correlating them to one another; 1000 iterations) was computed: *r*_*mean*_ = 0.50, *SE* = 0.03 (see Figure 3A).

**Figure 3.**
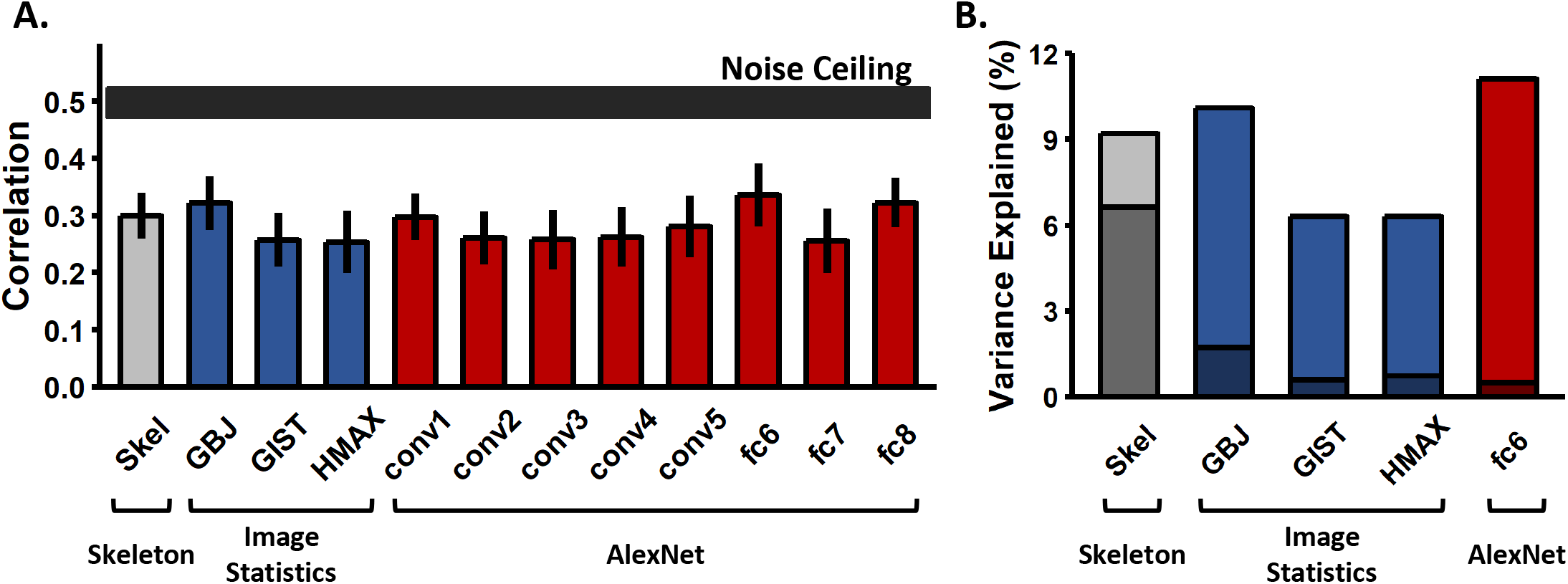
Results from Experiment 1. (A) Bar plot displaying the correlations (Pearson) between each model and human perceptual similarity judgments (error bars are bootstrapped SE). Models did not differ significantly from each other in the degree to which they predicted human judgments. The horizontal black bar represents the noise ceiling, which indicates the expected performance of the true model given the noise in the data (width represents SE). (B) Bar plot displaying the percentage of variance accounted for by each model individually (lighter shade), as well as the percentage of unique variance accounted for by each model (darker shade). A model of skeletal similarity explained the most unique variance (33.13% of total explainable variance) when compared to any single model or combination of models (see Supplemental Table 2 for the unique and shared variance explained by all model combinations).

Because the different models were predictive of participants’ judgments to similar degrees, and because objects with similar skeletons might also have similar image-level properties, it was important to test whether the different models accounted for the same variance in participants’ judgments, or whether a model of skeletal similarity explained unique variance. To this end, we conducted a regression analysis wherein all of the models and the most predictive layer of AlexNet (Skeleton ∪ GBJ ∪ GIST ∪ HMAX ∪ AlexNet-fc6) were included as predictors of human dissimilarity judgments. Together, these models explained 20.5% of the variance in human judgments, with Skeletal and GBJ models each explaining significant unique variance (*ps* < .01; see Supplemental Table 1). To ensure that the predictive power of the skeletal model was not simply the result of a suppression effect, we tested the skeletal model individually against every other model (Skeleton ∪ GBJ; Skeleton ∪ GIST; Skeleton ∪ HMAX; Skeleton ∪ AlexNet-fc6). Skeletal similarity was predictive of human judgments in each case (*r*^2^ = 14% –18%, *ps* < .001).

Finally, to better understand how the amount of variance explained by the skeletal model compared to the other models, we used variance partitioning analyses^50,51^. These analyses allowed us to determine how much of the total explained variance was unique to the different models and how much was shared by a combination of models. These analyses revealed that the model of skeletal similarity accounted for the greatest amount of unique variance in participants’ responses (6.6%) explaining 33.13% of the total explainable variance (see Figure 3B and Supplemental Table 2). A 4-predictor model consisting of the GBJ, GIST, HMAX, and AlexNet-fc6 models accounted for the next greatest amount of variance in participants’ responses (3.1%) accounting for 15.49% of the total explainable variance (see Supplemental Table 2). Taken together, these analyses suggest that, although other models of visual similarity were predictive of participants’ perceptual judgments, a model of skeletal similarity explained these judgments. These results suggest that skeletal structures may be an important source of information in making object identity.

A potential concern with these findings is that, because we created objects that varied in skeletal similarity, it was inevitable that a model of skeletal similarity would predict participants’ performance. We would emphasize, however, that other, non-skeletal models were also predictive of participants’ judgments, suggesting that participants incorporated other visual properties into their judgments. Nevertheless, to address this concern more directly, we tested whether the objects differed sufficiently for non-skeletal models to discriminate between them. A feature vector was extracted for every image (30 skeletons × 5 surface forms × 3 orientations) from each of these models (GBJ, GIST, HMAX, AlexNet-fc8). Then, for each model and object pair (same surface form), a linear support vector machine (SVM) classifier was trained to label objects using two object orientations; its ability to label the objects was tested using the third orientation. This procedure was repeated for every surface form and every combination of orientations between objects (0º × 0º; 0º × 30º; 0º × −30º; 30º × 30º; 30º × −30º; −30º × −30º). A final discrimination score was computed for each object pair by averaging the decoding accuracies across every surface form and combination of orientations. This analysis revealed that every model could discriminate between objects significantly above chance (0.50; *Ms* > 0.75), *ts* > 41.88, *ps* < .001, *ds* > 2.01 (see Supplemental Figure 1). Together, these findings demonstrate that the objects within our stimulus set were sufficiently different along other visual dimensions that non-skeletal models could accurately discriminate them.

### Experiment 2 – Can perceived similarity be explained by another model of structure?

The results of Experiment 1 suggest that humans incorporate skeletal representations when making object similarity judgments. However, an alternative possibility is that participants’ sensitivity reflected a different model of structure, namely one based on the coarse spatial relations between object parts^52–54^. A model based on coarse spatial relations suggests that the structure of an object is represented by the categorical relations between component parts (e.g., components above one another vs. components side-by-side). In contrast to skeletal descriptions of shape, which describe quantitative relations between component parts, a coarse spatial-relations model would predict that only qualitative changes to the overall spatial arrangement of the parts (e.g., changing component relations from ‘side-by-side’ to ‘end-to-end’) should influence object recognition. Yet objects with similar spatial relations also have more similar skeletal structures. Thus, the relation between skeletal similarity and human perceptual judgments in Experiment 1 could have reflected the co-variation between the shape skeleton and an object’s coarse spatial relations.

Here we tested whether participants’ judgments of perceptual similarity were influenced by an object’s skeletal structure even when coarse spatial relations were held constant, and thus unable to be used as a similarity cue. If perception of object shape is based on a skeletal structure, then a proportional change to the shape skeleton would elicit a proportional decrease in recognition, even when the coarse spatial relations are unchanged. Thus, as two skeletons become more dissimilar, participants should judge the objects as more different from one another.

#### Stimuli and experimental design

To test whether proportional changes to the shape skeleton led to proportional deficits in recognition, we adapted three objects from Experiment 1 to have six increments of skeletal dissimilarity (0%, 10%, 20%, 30%, 40%, 50% difference; see Methods for additional details). The three objects consisted of distinct coarse spatial relations (see Figure 4A, Methods, and Supplemental Figure 2). Changes to the skeleton were implemented by moving one component along the length of another component in 10% increments (see Figure 4A). This manipulation caused systematic changes to the shape skeleton without changing the coarse spatial relations between the object’s component parts. Thus, if skeletal similarity affects participants’ perceptual judgments, independent of coarse spatial relations, then performance should scale proportionally with changes to the skeleton.

**Figure 4.**
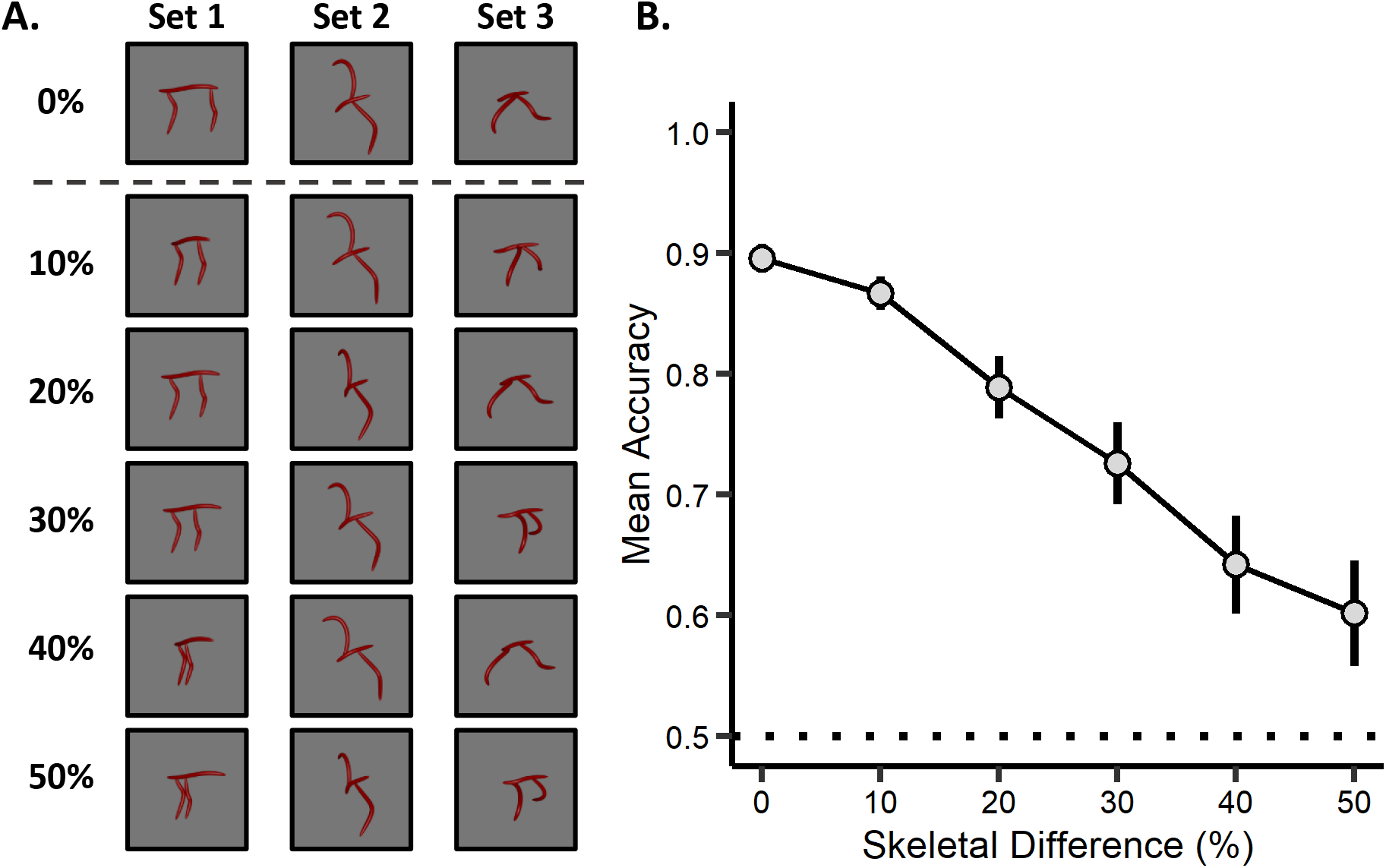
Example stimuli and results from Experiment 2. (A) Objects were comprised of three sets, each with distinct coarse spatial relations (separate columns). Crucially, objects with the same spatial relations varied in skeletal similarity by increments of 0%, 10%, 20%, 30%, 40%, or 50% (each row within a column). On the ‘same’ test trials (objects within the same column), participants were presented with a reference object (0%; top row) and an object with the same coarse spatial relations. On the ‘different’ test trials (objects across columns), participants were presented with objects that had different coarse spatial relations. Objects were presented in one of three orientations (30°, 60°, 90°; see Supplemental Figure 2 for full stimulus set). (B) Participants’ recognition accuracy (proportion correct) for objects with the same coarse spatial relations decreased as a function of skeletal change, suggesting that humans represent object structure by their skeletons. The dotted line represents chance performance and the error bars represent SE.

Participants (*n* = 42) completed a discrimination task in which they were shown two simultaneously presented objects and were instructed to judge whether the objects were the same or different in their coarse spatial relations. Crucially, participants were instructed to ignore any changes to the precise positions of the object parts (i.e., exact skeleton) so as to make their decision on the basis of “overall shape.” Participants were given a familiarization phase to ensure they understood that these instructions referred to objects with the same coarse spatial relations (e.g., in Figure 4A each column consists of objects with the same “overall shape”, but different skeletons). Objects were presented from three orientations that maximized the visibility of the object’s structure (30°, 60°, and 90°). Participants were given unlimited time to respond but were encouraged to respond quickly and accurately.

#### Results and discussion

Participants performed significantly above chance (0.50) at every level of skeletal change, *Ms* > 0.60, *ts*(34) > 2.32, *ps* < 0.026, *ds* > 0.39, (*M*_*RTs*_ < 1705 ms), demonstrating that they followed the task instructions to identify objects by their coarse spatial relations. Crucially, however, participants’ performance in discriminating between objects with the same coarse spatial relations was less accurate as a function of skeletal change, *F*(1, 34) = 51.77, *p* < 0.001, η_p_^2^ = 0.60 (Figure 4B), suggesting that when participants make perceptual similarity judgments, they incorporate fine-grained structural information, as predicted by a skeletal model. Nevertheless, as in the previous experiment, a change to the object’s shape skeleton also induced changes along other visual dimensions (e.g., image-statistics). Thus, we tested whether participants’ performance was better described by other models of vision. To this end, we used a random-effects regression analysis, with the skeletal similarity model and other models (i.e., GBJ, GIST, HMAX, and AlexNet-fc6) as predictors (subject and object as the random effects; see Methods for additional details). Analyses revealed that the model of skeletal similarity remained a significant predictor of human performance, even when controlling for the other models, χ^2^(1) = 22.30, *p* < 0.001, and that it explained the greatest amount of variance in participants’ responses, β = −1.24. The only other model to explain unique variance was GIST, χ^2^(1) = 19.55, *p* < 0.001, β = 0.63, suggesting a role for image-statistics in this process (see Supplemental Table 3 for the results of the other models). To ensure that the predictive power of the skeletal model was not simply the result of a suppression effect, we tested the skeletal model against every other model (Skeleton ∪ GBJ; Skeleton ∪ GIST; Skeleton ∪ HMAX; Skeleton ∪ AlexNet-fc6). Skeletal similarity was predictive of human judgments in each case, χ^2^(1) > 9.27, *ps* < 0.002. Taken together, these findings suggest that participants’ judgments of perceived object similarity reflect the metric positions of object parts, consistent with an object representation based on skeletal structure, not the course spatial relations. Combined with Experiment 1, these results provide further support for the unique, and possibly privileged, role of skeletal descriptions of shape in object recognition.

### Experiment 3 – Are skeletal structures a privileged source of information for object recognition?

A model of skeletal similarity was most predictive of human perception in Experiments 1 and 2, when compared to other, non-shape-based, models of visual similarity (i.e., GBJ, GIST, HMAX, and AlexNet), as well as another descriptor of structure (i.e., coarse spatial relations). Together, these results suggest that shape information, particularly skeletal descriptors of shape, plays an important role in object recognition. However, there exist alternative descriptors of shape, which emphasize local contour information^55,56^ or the non-accidental properties (NAPs) of component parts^12^, not skeletal structures. Thus, it remains unknown whether, for object recognition, skeletal structures offer a more informative descriptor of shape than local contour information and component parts.

To test this hypothesis, the skeleton of an object was pitted against its surface form in a match-to-sample task. Surface forms were designed to alter the object’s contours without changing the object’s underlying skeleton^37^ (see Figure 2B). As described in more detail below, surface form similarity was perceptually matched to skeletal similarity and surface forms were well characterized by other models of vision. Moreover, surface forms were created such that they differed in NAPs in order to compare the skeletal descriptions against a model of shape based on component parts^12,57^. NAPs, such as the degree to which a component tapers or bulges outward, are thought to play an essential role in models of shape perception because they serve as unique identifiers of component parts, allowing objects to be identified from a variety of viewpoints^12,58^. Thus, by pitting an object’s skeleton against its surface form, we can better understand the degree to which different descriptors of shape are used for object recognition.

#### Surface form properties

To quantify the degree of visual similarity between surface forms, participants (*n* = 41) conducted a surface form discrimination task (see Methods). In this task, participants judged whether two objects were the same or different in surface form (same skeletons). Surface form discrimination accuracy was compared to skeletal discrimination accuracy from Experiment 1 for the four skeletons used here (see Methods and Supplemental Figure 3). This analysis revealed that surface form discrimination accuracy (*M* = 0.87, *SD* = 0.19) did not differ from skeleton discrimination accuracy (*M* = 0.86, *SD* = 0.20), *t*(77) = 0.76, *p* = 0.94. Follow up analyses revealed that participants’ surface form discrimination performance was well described by GBJ, GIST, HMAX, and AlexNet-fc6 models, *rs* = 0.63-0.77, with AlexNet-fc6 explaining unique variance when all four models were entered into a random-effects regression, χ^2^(1) = 12.71, *p* < 0.001.

To test whether surface forms were comprised of unique NAPs, a separate group of participants (*n* = 41) were taught to identify different NAPs and they then rated the degree to which each surface form exhibited a particular NAP (e.g., “To what extent do parts of this object exhibit taper?”) on a 7-point Likert scale (1 “not at all”; 7 “a lot”; see Methods for details). A *k*-means cluster analysis^59^ revealed that participants’ ratings were best described by four clusters, and a permutation test, in which cluster labels were shuffled 10,000 times, revealed that cluster labels were predictive of each surface form significantly better than chance (*ps* < 0.002). That the surface forms were better described by four, rather than five (one for each surface form), clusters is consistent with two of the surface forms having the same NAPs, but differing in metric properties such as circumference (see Figure 3B)^57,60^.

#### Match-to-sample task: design

Having confirmed that the surface forms were perceptually matched to skeletal differences, and that they were comprised of unique NAPs, we were in a position to test whether skeletal descriptions of shape were a privileged source of information for object recognition relative to other descriptors of shape. In a match-to-sample task, a separate group of participants (*n* = 39) were presented with a sample object and two choice objects (i.e., target and distractor; see Figure 5A-C). They were instructed to judge which of the two choice objects was most likely to be from the same category as the sample object. The target object matched the sample object in either its skeleton or surface form. The distractor object differed from the sample object by both skeleton and surface form. These trials ensured that participants were able to match objects by either their skeleton or surface form when each property was presented in isolation (see Supplemental Figure 5A-B). Other trials presented a conflict between the object’s skeleton and surface form such that one of the choice objects matched the sample’s skeleton, but not surface form, and the other object matched the object’s surface form, but not skeleton (see Figure 5C). The conflict trials tested whether the skeletal descriptors served as a preferred cue for object recognition. The objects were presented as still images in one of three depth orientations (30°, 60°, 90°; see Supplemental Figure 3). Participants were instructed to ignore the orientations of the objects and, on each trial, to choose which of the two choice objects was from the same category as the sample object.

**Figure 5.**
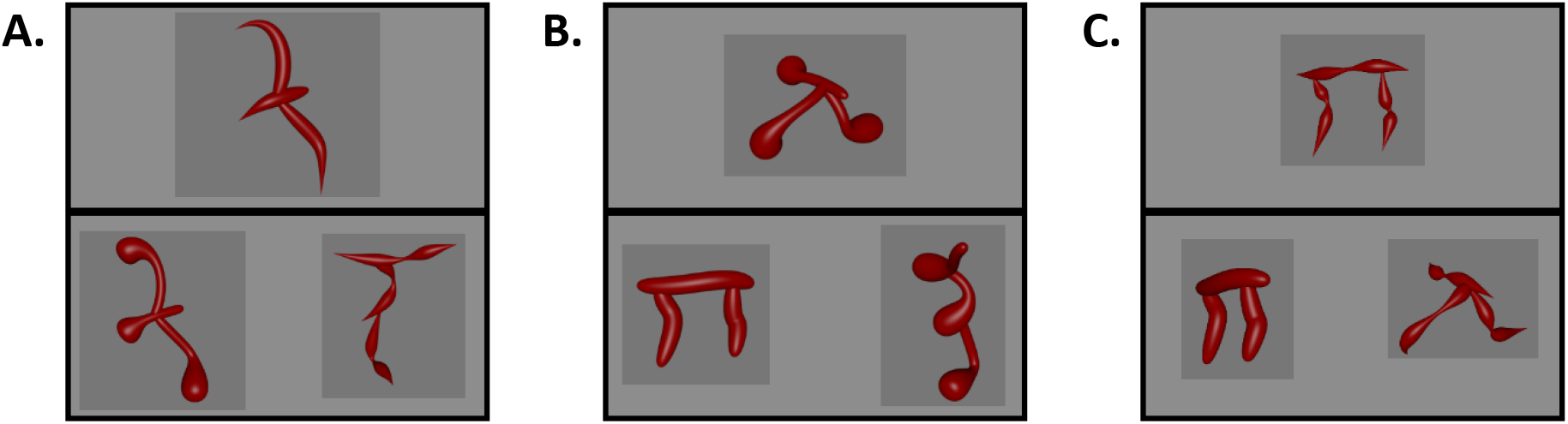
Examples of the three trial types used in Experiment 3. (A) A skeleton match trial wherein one choice object matched the sample’s skeleton, but not surface form. The other choice object matched on neither skeleton nor surface form. (B) A surface form match trial wherein one choice object matched the sample’s surface form, but not skeleton. The other choice object matched on neither skeleton nor surface form. (C) A conflict trial wherein one choice object matched the sample’s skeleton, but not surface form, and the other choice object matched the sample’s surface form, but not skeleton.

#### Match-to-sample task: results and discussion

Participants successfully categorized objects by either their skeletons, *M* = 0.88 (*M*_*RT*_ = 1167 ms), *t*(38) = 27.01, *p* < 0.001, *d* = 4.32, 95% CI [3.35, 5.41], or surface forms, *M* = 0.78 (*M*_*RT*_ = 1419 ms), *ts*(38) = 15.51, *p* < 0.001, *d* = 2.48, 95% CI [1.87, 3.16], when each cue was presented in isolation, as indicated by their above chance performance in these conditions (see Figure 6A). Crucially, however, on the conflict trials, participants categorized objects by their skeletons, not surface forms, *t*(38) = 6.63, *p* < 0.001, *d* = 1.06, 95% CI [0.66, 1.45] (see Figure 6B-C). Indeed, participants preferentially categorized objects by their skeletons when pitted against all, *M* > 0.61 (*M*_*RT*_ < 1218 ms), *t*s[38] > 2.52, *ps* < .016, *ds* > 0.40) but one, *M* = 0.58 (*M*_*RT*_ = 1140 ms) (*p* = .059, *d* = 0.31) surface form. Thus, although surface forms were perceptually matched to the objects’ skeletons and were comprised of unique NAPs, participants relied more heavily on the shape skeleton when classifying objects, suggesting that skeletal structure may be a privileged source of shape information for object recognition.

**Figure 6.**
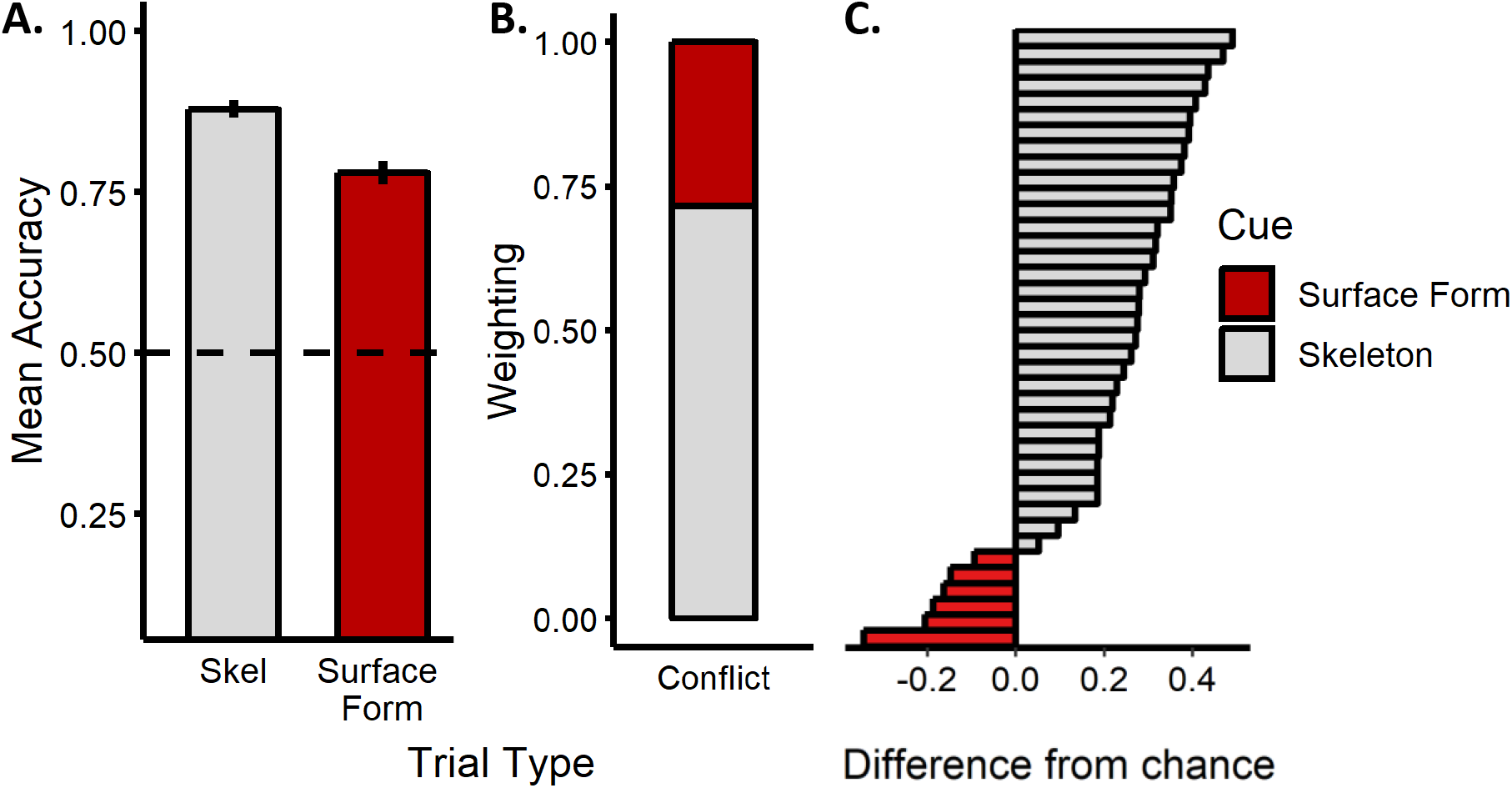
Results from the match-to-sample task of Experiment 3. (A) Participants’ mean accuracy (error bars represent SE) on trials in which only a skeleton or surface form match was possible (dotted line indicates chance performance). (B) Participants’ categorization judgments in the conflict trial. A value closer to 1 indicates greater weighting of the object’s skeleton; a value closer to 0 indicates greater weighting of the object’s surface form. Although participants successfully matched objects by their skeletal structure or surface forms when each cue was presented in isolation, they were more likely to match objects by their skeleton, as opposed to their surface forms, when these cues conflicted with one another. (C) Histogram of participants’ responses on the conflict trials. A value greater than zero indicates greater weighting of skeletal information. The majority of participants matched objects by their skeleton, demonstrating a consistent pattern of responses across participants.

## General Discussion

The ability to determine the similarity between shapes is crucial for object recognition. Shape skeletons may be particularly useful in this context because they provide a compact descriptor of shape, as well as a formalized method for computing shape similarity. Nevertheless, few models of biological object recognition include skeletal descriptions in their implementation. Here we tested whether skeletal structures provide an important source of information for object recognition when compared with other models of vision. Our results showed that a model of skeletal similarity was most predictive of human object judgments when contrasted with models based on image-statistics or neural networks, as well another model of structure based on coarse spatial relations. Moreover, we found that skeletal structures were a privileged source of information when compared to other properties thought to be important for shape perception, such as object contours and component parts. Thus, our results suggest that not only does the visual system show sensitivity to the skeletal structure of objects^32,36,37^, but also that perception and comparison of object skeletons may be crucial for successful object recognition.

The strength of skeletal models is that they provide a compact description of an object’s global shape structure, as well as a metric by which to determine shape similarity. Indeed, shape skeletons may offer a concrete formalization for the oft poorly defined concept of global shape. Skeletal descriptions exhibit many properties ascribed to global shape percepts such as relative invariance to local contour variations^22,35^. Moreover, there exist many methods by which to compare skeletal structures, such as by their hierarchical organization^61^ or using distance metrics (as used here)^27^, thereby allowing for a quantitative description of shape similarity. Such a description may be particularly important when recognizing objects across previously unseen views or categorizing novel object exemplars.

In the current study, we chose to test whether humans represent a 3D skeletal structure rather than a 2D skeleton that is arguably more easy to compute from an image^26^. Our decision was motivated by behavioral^36^ and neuroimaging work^37,38^ suggesting sensitivity to 3D skeletons in the primate visual system, as well as accumulating evidence that object perception (at least for novel objects) is best described by a 3D object-centered shape representation^48,49^. However, it remains unknown how a 3D skeletal structure arises from 2D images on the retina. One possibility is that skeletal computations in the visual system invoke generative shape processes^62,63^. These processes may be able to recover an object’s 3D skeletal structure from retinal images by incorporating a small number of image-computable 2D skeletons (e.g., one from each eye)^64^. Alternatively, it is possible that an object’s 3D structure may be recovered from a single image by first creating a representation based on depth properties and surface orientation, a so-called 2.5D sketch^8^. Indeed, recent neural network models have been able to successfully reconstruct an object’s 3D shape from single images by incorporating 2.5D sketches^65^, offering a possible mechanism for 3D skeleton generation. Consistent with these possibilities, our results showed that object judgments were well described by a 3D skeletal structure even though participants were only ever shown object images from a limited number of viewpoints. Nevertheless, more research is needed to understand how 3D skeletal representations arise from 2D images on the retina.

A question that arises from the current findings is the extent to which the stimuli and tasks used here invoke the same mechanisms as rapid real-world object perception, also known as ‘core’ object recognition^66^. Indeed, one might ask whether the tightly controlled stimulus set, which was designed to vary in skeletal similarity, and the untimed tasks, where participants could directly compare the similarity of objects, invoke ‘core’ object recognition processes. It is well known that the visual system receives input from multiple systems (e.g., frontal and parietal regions) and incorporates recurrent processes to solve object recognition, particularly in cases of uncertainty^67–69^. Thus, it is possible that object recognition tasks in this study, and the implementation of skeletal models more generally, may have invoked higher-level processes. Although we acknowledge that object recognition is not a unitary process, with higher-level processes playing an important role, we suggest that our stimuli and tasks likely measured core object recognition. In particular, in Experiment 1, we found that the objects could be discriminated by models that are implemented during a feedforward sweep through the ventral visual stream, and these models were also predictive of human judgments. Moreover, we found that participants performed equally well when objects were presented for only 100 ms (see Supplemental Experiment 1). Nevertheless, it is an open question whether shape skeletons are implemented using exclusively feedforward mechanisms or whether recurrent or generative processes are also needed^6,63,70^.

Although our findings suggest that skeletal descriptors play an important role in object recognition, we would not argue that skeletal descriptions alone are sufficient. Humans do not perceive the visual environment simply as a collection of skeletons, but rather, as complete objects where local contours, textures, and colors are integrated with a shape structure. Indeed, our results showed that other models of vision were also predictive of participants’ object judgments to varying degrees in every experiment. That other models were also predictive may be unsurprising, given that both shape- and non-shape-related properties are known to play important roles in object recognition. For instance, local contour information may be particularly useful for making subordinate-level category distinctions where the skeletons of objects are roughly the same^54,71^. Similarly, texture statistics and feature descriptions have been shown to be important indicators of both basic^11,72^ and superordinate-level^73,74^ object distinctions. And though it is currently unclear whether feedforward neural network models (such as the one tested here) incorporate global shape information^75,76^, more biologically plausible models with recurrent or generative architectures may begin to approximate human shape perception^68^. Nevertheless, our work highlights the importance of formalized models of shape for object recognition, particularly the unique, and possibly privileged, role that skeletal structures may play.

## Methods

### Participants

205 participants were tested in the current study (*M*_age_ = 19.75 years; *range* = 18.03 – 23.48 years). Of these participants, 16 were excluded for exhibiting chance, or below chance, performance (3 from Experiment 1; 7 from Experiment 2; 3 from the surface form discrimination task of Experiment 3). Because accuracy could not be evaluated in the NAP rating task of Experiment 3, we ensured that participants exhibited reliable performance; 6 participants were excluded from this experiment for failing to meet this criterion (αs < 0.7). All participants provided informed consent and participated for course credit. Experimental procedures were approved by Emory University’s Institutional Review Board (IRB). All experiments were performed in accordance with the relevant guidelines and regulations of the IRB.

### Apparatus

All tasks were presented on a desktop computer with a 19-inch screen (1280 × 1024 px) and controlled using custom software written in Visual Basic (Microsoft). Participants sat at a distance of ~60 cm from the computer screen.

### Experiment 1

#### Stimulus generation

Objects were procedurally generated using the Python API for Blender (Blender Foundation). Each skeleton was comprised of three segments created from Bezier curves of a random size and curvature scaled between .05 and .25 virtual Blender units (vu). The first axis segment was oriented forward towards the ‘camera’. The second and third segments were oriented perpendicular to the first segment and attached to the first segment or second segment at a random point along their length. Surface forms were created by applying a circular bevel to the object’s skeleton along with one of five taper properties that determined the shape of the surface form. Finally, the overall size of the object was normalized to. 25 vu.

#### Skeletal similarity model

The coordinates of the skeleton for each object were extracted by sampling 999 points along the length of each axis segment (2997 points in total). Skeletal points were normalized by the length of each segment by subsampling points along the skeletal structure until these points were evenly spaced across the skeleton by 0.0005 vu (for ease of analysis, coordinates were rescaled by a factor of 1000). Skeletal similarity was calculated as the mean Euclidean distance between each point on one skeleton structure with the closest point on the second skeleton structure following maximal alignment. Maximal alignment was achieved by overlaying each structure by its center of mass and then iteratively rotating each object in the picture plane orientation by 15° until the smallest distance between structures was found.

#### Gabor-jet (GBJ) model

The GBJ model is a low-level model of image similarity inspired by the response profile of complex cells in early visual cortex^46^. It has been shown to scale with human psychophysical dissimilarity judgments of faces and simple objects^77^. To simulate the response profile of complex cell responses, the model overlays a 12 × 12 grid of Gabor filters (5 scales × 8 orientations) along the image. The image is convolved with each filter, and the magnitude and phase of the filtered image is stored as a feature vector. Dissimilarity between each image is computed as the mean Euclidean distance between feature vectors of each image. A single dissimilarity value was computed for each object pair by taking the mean Gabor activation distance for an object pair across the three orientations (30 objects × 3 orientations).

#### GIST

The GIST model is considered a mid-level model of image similarity that describes the content of an image through global image features^47^. It has been shown to accurately describe the content of natural images, particularly as they relate to scene perception^39^. The model overlays a grid of Gabor filters (4 scales × 8 orientations) on the image and then convolves the image with the filters, creating a feature activation map. This feature map is divided into 16 regions (based on the 4 × 4 grid) and then mean activation within each region is computed and stored as a GIST feature vector. GIST dissimilarity between each image is computed as the mean Euclidean distance between feature vectors of each image. A single dissimilarity value was computed for each object pair by taking the mean Euclidean distance for an object pair across the three orientations (30 objects × 3 orientations).

#### HMAX

The HMAX (hierarchical MAX) model is a biologically inspired hierarchical neural network model that describes an image by max-pooling over a series of simple (S1, S2) and complex (C1, C2) units^40,78^. It has been shown to match human performance on simple category judgment tasks (e.g., animals and non-animals) and exhibits some invariance to changes in position and scale. In the current study, we used the feature patches provided with the HMAX model. In the first layer (S1), each image is convolved with Gabor filters (8 scales × 4 orientations), the output of which is fed into a second layer (C1) that determines the local maximum over all positions and scales. The outputs of these layers are fed through a second set of simple and complex units (S2, C2). Dissimilarity was computed by extracting the C2 activations for each image and correlating (Pearson) it with the activations for every other image. A single dissimilarity value was computed for each object pair by taking the mean correlation for an object pair across the three orientations (30 objects × 3 orientations).

#### CNN

As a model of high-level vision, we used AlexNet, an eight layer CNN pre-trained to classify objects from the ImageNet database^41,79^. We adopted AlexNet rather than other CNNs in our analyses because its architecture is relatively simple by comparison and because it can identify objects with high accuracy^41^. Importantly, AlexNet has been shown to be predictive of human object similarity judgments^42^. CNN similarity for each object was calculated by extracting a feature vector from each convolutional, and fully connected, layer for each object image, and then computing the mean Euclidean distance between the feature vector for each image with every other image. A single difference value was computed for each object pair by taking the mean CNN difference for an object pair across the three orientations (30 objects × 3 orientations). Dissimilarity values were calculated for every layer and correlated to participants’ behavioral judgments. Participants’ similarity judgments in Experiment 1 were most strongly correlated with fully-connected layer 6 (fc6) of AlexNet, *r* = 0.34 (see Figure 3A). Because fc6 was most predictive in this experiment, we tested only fc6 in subsequent experiments.

#### Discrimination task

On each trial, participants were shown images of two objects (side-by-side) presented simultaneously in one of three depth orientations (−30°, 0°, +30°). Objects were matched for surface form and either had the same or different skeleton. Participants were instructed to decide whether the two images showed the same or different object (independent of orientation). Each object was paired with every other object (including itself) during the experimental session. Participants were administered a total of 885 trials (435 different and 450 same trials). Each trial began with a fixation cross (500 ms), followed by a pair of objects, which remained onscreen until a response was made, followed by an inter-trial interval (500 ms). Each object was approximately 6°× 6° in size and subtended 9° from the center of the screen.

### Experiment 2

#### Stimulus generation and model analyses

A *k*-means cluster analysis was conducted on participants’ discrimination data from Experiment 1. This analysis revealed that objects were well described by four clusters. Based on these clusters, three perceptually matched objects were chosen whose skeletons could be altered without changing the coarse spatial relations (see Supplemental Figure 2). Importantly, these objects also had different coarse spatial relations (Set 1: two components below a third, pointing down, one component placed in front of the other; Set 2: one component on either side of a third, one pointing up and the other down; Set 3: one component on either side of a third, each pointing diagonally down). Six versions of each object (0%, 10%, 20%, 30%, 40%, and 50% skeleton difference) were generated by moving one segment along the length of the central segment in 10% increments (relative to the length of the central component). Objects were rendered with only the thinnest surface form to prevent component parts from overlapping. Images of each object (18 total) were generated in three orientations (30°, 60°, 90°) intended to maximize the view of each object. Each object was analyzed and compared with every other object using the same models and procedure described in Experiment 1.

#### Discrimination task

On each trial, participants were shown images of two objects (side-by-side) presented simultaneously in one of the three depth orientations (30°, 60°, 90°). Each object was rendered with the same surface form (see Figure 4A) and either had the same or different coarse spatial relations. On each ‘same’ trial, participants were presented with both a reference object (0% skeletal difference) and another object that had the same coarse spatial relations but different skeleton in one of the increments described previously (objects in the same columns of Figure 4A). On each ‘different’ trial, participants were presented with two objects that had different coarse spatial relations (any possible skeleton; objects in the same rows of Figure 4A). Participants were instructed to decide whether the two images showed an object with the same or different “overall shape” (independent of orientation). Participants were given instructions and 8 sample trials (with feedback) using a separate set of objects to ensure that they understood that “overall shape” referred to objects with the same coarse spatial relations (4 same trials; 4 different trials). In the same trials, each skeletal difference was presented an equal number of times in each possible orientation. In the different trials, object pairs with different coarse spatial relations (any possible skeleton) were randomly selected and presented in randomly determined orientations. Participants were administered at total of 648 trials (324 same trials and 324 different trials). Each trial began with a fixation cross (500 ms), followed by a pair of objects, which remained onscreen until a response was made, followed by an inter-trial interval (500 ms). Each object was approximately 6° × 6° in size and subtended 9° from the center of the screen.

### Experiment 3

#### Stimulus generation and model analyses

Four perceptually matched objects were chosen from the object clusters identified in Experiment 2. Images of each object (4 objects × 5 surface forms) were generated in three orientations (30°, 60°, 90°) intended to maximize the view of each object (see Supplemental Figure 2).

#### Surface form discrimination task

On each trial, participants were shown images of two objects (side-by-side) in one of the three depth orientations (30°, 60°, 90°). Objects had the same shape skeleton and either the same or different surface forms. Participants were instructed to decide whether the two images showed the same or different object (independent of orientation). Each surface form was paired with every other surface form an equal number of times for a total of 600 trials.

#### NAP task

To test whether surface forms were comprised of unique component parts, participants rated each surface form on the degree to which it exhibited a specific NAP. During a training phase, participants were taught a subset of NAPs (drawn from Amir, et al. ^57^) and then shown a subset of objects that they were asked to rate on the degree to which they exhibited the specific NAP. The four NAPs were: (1) *taper*, defined as the degree to which the thickness of an object was reduced towards the end (taper in the current study corresponds to ‘expansion of cross-section’ in Amir et al.^57^); (2) *positive curvature*, defined as the degree to which an object part curved outwards; (3) *negative curvature*, defined as the degree to which an object part curved inwards; and (4) *convergence to vertex*, defined as the degree to which an object part ended in a point. We excluded the curved versus straight axis property of Amir et al.^57^ because it was confounded with the object’s skeleton. We also excluded the change in cross-section property (e.g., circular vs. rectangular shape) because all of the surface forms had a circular cross-section. Participants were tested on their understanding of the four NAPs with a task in which they were presented with pairs of single-part objects (simultaneously onscreen) where one exhibited an NAP and the other did not. Participants were instructed to select the object that exhibited more/less of a particular NAP (e.g., “Which object exhibits more positive curvature?”; feedback was provided). During the rating phase, participants were shown each test stimulus with each surface form (4 objects × 5 surface forms) and asked to rate the degree to which each surface form exhibited a particular NAP (e.g., “To what extent do parts of this object exhibit taper?”) on a 7-point Likert scale (1 = “not at all”; 7 = “a lot”).

#### Match-to-sample task

On each trial, participants were shown one object (sample) placed centrally near the top of the screen above two objects near the bottom of the screen (target and distractor). Participants were instructed to choose which of the two bottom objects was most likely to be in the same category as the sample. Participants were presented with three possible trial types: skeleton and surface form trials, in which one object matched the sample in either skeleton or surface form, respectively (the other object matched on neither; see Figure 5A-B); and conflict trials in which one object matched in skeleton, but not surface form, and the other object matched in surface form, but not skeleton (see Figure 5C). Participants were administered a total of 480 trials (160 of each trial type). Each trial began with a fixation cross (500 ms), followed by the sample and choice objects, which remained onscreen until a response was made, followed by an inter-trial interval (500 ms). Each stimulus was approximately 6° × 6° in size, and choice objects subtended 9° from the center of the screen.

## Supplemental Materials

### Supplemental Experiment 1

One potential concern with our stimuli is that they may have been difficult to discriminate because they may represent a single class of unfamiliar objects with a high degree of visual similarity. Such objects may be less likely to elicit the same mechanisms that support ‘core’ object recognition, which is thought to occur within 100-200 ms via a feedforward sweep through the ventral stream. Instead, discrimination of these objects may require additional high-level processes, such as mental rotation^80^, not typically implemented when discriminating familiar objects. This possibility is difficult to rule out in the main experiments because participants were given unlimited time to discriminate between objects. Thus, in a supplemental experiment, we tested participants in a speeded task where the target object was presented for a 100 ms. If performance on this speeded task were comparable to performance on the unspeeded task (see Experiment 1 in the main text), then it would suggest that both tasks could measure core object recognition.

Participants (*n* = 14) were administered a sequential match-to-sample task where they were asked to decide which of two choice objects matched a previously presented sample object. Each trial began with a fixation cross (500 ms), followed by a display with the sample object (100 ms), and then a display with two choice objects which remained onscreen until a response was made. One choice object had the same skeleton as the sample, and the other choice object had a different skeleton. The choice objects always had the same surface form as the sample (randomly selected) but were presented from different orientations (−30°, 0°, 30°). Participants were instructed to ignore the orientations of the objects and to make their decision on the basis of visual similarity. Each object was pitted against every other object an equal number of times (435 trials). Each object was approximately 6° × 6° in size, and choice objects subtended 9° from the center of the screen.

Comparisons to chance (0.50) revealed that participants were able to match the sample object with the correct choice object, *M* = 0.82% (*M*_*RT*_ = 946 ms), *t*(13) = 26.8, *p* < .001, *d* = 7.16, with 14/14 participants displaying accuracy above 0.74. This result suggests that our objects differed sufficiently to allow object recognition to occur within 100 ms.

In a subsequent analysis, we tested whether participants’ performance on this task differed from their performance in the unspeeded discrimination task used in Experiment 1. We found that participants’ performed comparably in the two tasks (*M*_accuracy_ = 0.82 vs. 0.80), with no statistical difference between groups (*p* = 0.46). These results are consistent with the speeded and unspeeded tasks recruiting similar perceptual processes, namely ‘core’ object recognition.

---

**Supplemental Table 1.** Linear regression results for each model used in Experiment 1.

| Model | Standardized<br>Coefficient | *t* | *p* |
| --- | --- | --- | --- |
| | $\beta$ | | |
| (Constant) |  | 1.82 |  |
| Skeleton | 0.27 | 6.01 | < .001 |
| Gabor-Jet | 0.31 | 3.17 | 0.002 |
| GIST | -0.18 | -2.03 | 0.043 |
| HMAX | 0.11 | 2.06 | 0.040 |
| AlexNet-fc6 | 0.13 | 1.80 | 0.073 |

**Supplemental Table 2.** Coefficients displaying the percentage of unique and shared variance explained by each model and model combinations, as well as percentages of the total explainable variance (20.5%) explained by each model and model combinations.

| Model | Coefficient | Percentage of total |
| --- | --- | --- |
| Skel | 6.631 | 33.305 |
| GBJ | 1.726 | 8.669 |
| GIST | 0.587 | 2.949 |
| HMAX | 0.732 | 3.678 |
| fc6 | 0.499 | 2.508 |
| Skel + GBJ | -0.641 | -3.218 |
| Skel + GIST | -0.241 | -1.209 |
| GBJ + GIST | -0.512 | -2.569 |
| Skel + HMAX | -0.171 | -0.859 |
| GBJ + HMAX | 0.344 | 1.730 |
| GIST + HMAX | 0.037 | 0.187 |
| Skel + fc6 | 1.005 | 5.046 |
| GBJ + fc6 | 1.218 | 6.116 |
| GIST + fc6 | -0.105 | -0.525 |
| HMAX + fc6 | 0.445 | 2.235 |
| Skel + GBJ + GIST | 0.224 | 1.125 |
| Skel + GBJ + HMAX | -0.095 | -0.477 |
| Skel + GIST + HMAX | -0.009 | -0.047 |
| GBJ + GIST + HMAX | 0.032 | 0.162 |
| Skel + GBJ + fc6 | 0.299 | 1.502 |
| Skel + GIST + fc6 | -0.047 | -0.237 |
| GBJ + GIST + fc6 | 1.852 | 9.302 |
| Skel + HMAX + fc6 | 0.236 | 1.187 |
| GBJ + HMAX + fc6 | 1.017 | 5.109 |
| GIST + HMAX + fc6 | -0.039 | -0.196 |
| Skel + GBJ + GIST + HMAX | -0.006 | -0.032 |
| Skel + GBJ + GIST + fc6 | 1.001 | 5.027 |
| Skel + GBJ + HMAX + fc6 | 0.037 | 0.186 |
| Skel + GIST + HMAX + fc6 | 0.009 | 0.045 |
| GBJ + GIST + HMAX + fc6 | 3.084 | 15.493 |
| Skel + GBJ + GIST + HMAX + fc6 | 0.758 | 3.809 |

**Supplemental Table 3.**
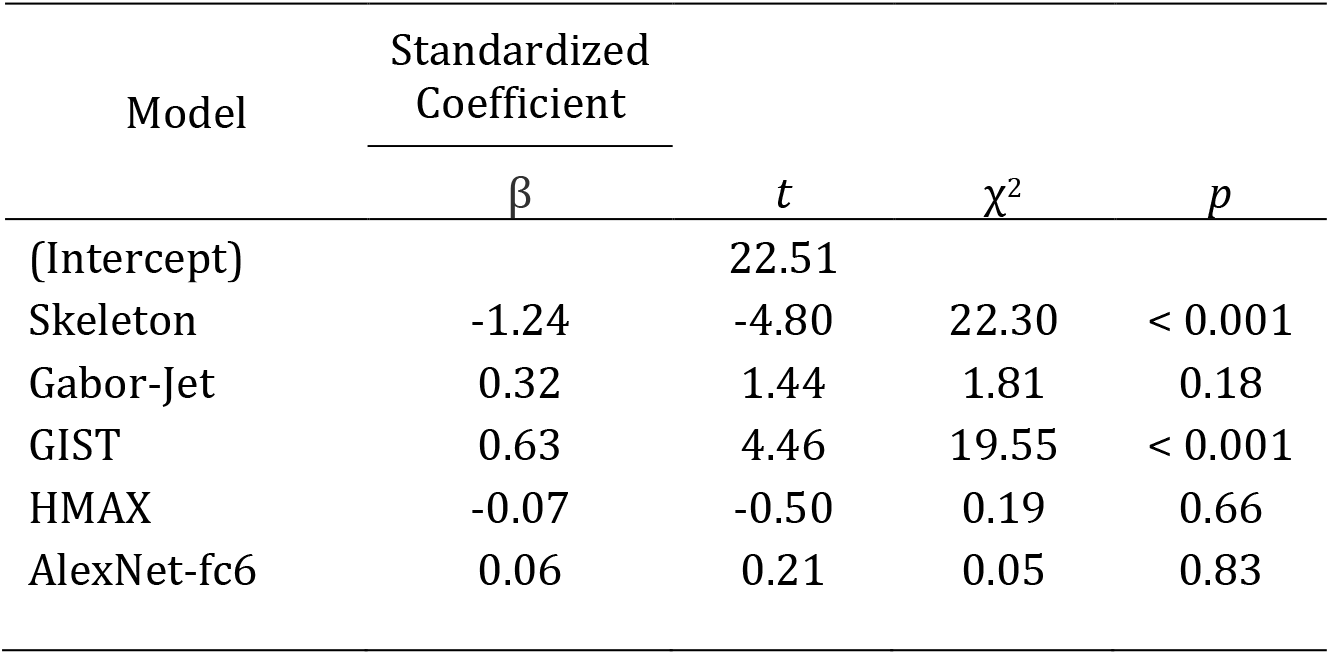
Random-effects regression results for each model used in Experiment 2. The standardized coefficients and *t*-values are drawn from a full regression model that included each model as a predictor. The χ^2^ and *p-*values are calculated by iteratively testing the full regression model against ones without the predictor of interest.

| Model | Standardized<br>Coefficient | $t$ | $\chi^2$ | $p$ |
| --- | --- | --- | --- | --- |
| | $\beta$ | | | |
| (Intercept) |  | 22.51 |  |  |
| Skeleton | -1.24 | -4.80 | 22.30 | < 0.001 |
| Gabor-Jet | 0.32 | 1.44 | 1.81 | 0.18 |
| GIST | 0.63 | 4.46 | 19.55 | < 0.001 |
| HMAX | -0.07 | -0.50 | 0.19 | 0.66 |
| AlexNet-fc6 | 0.06 | 0.21 | 0.05 | 0.83 |

**Supplemental Figure 1.**
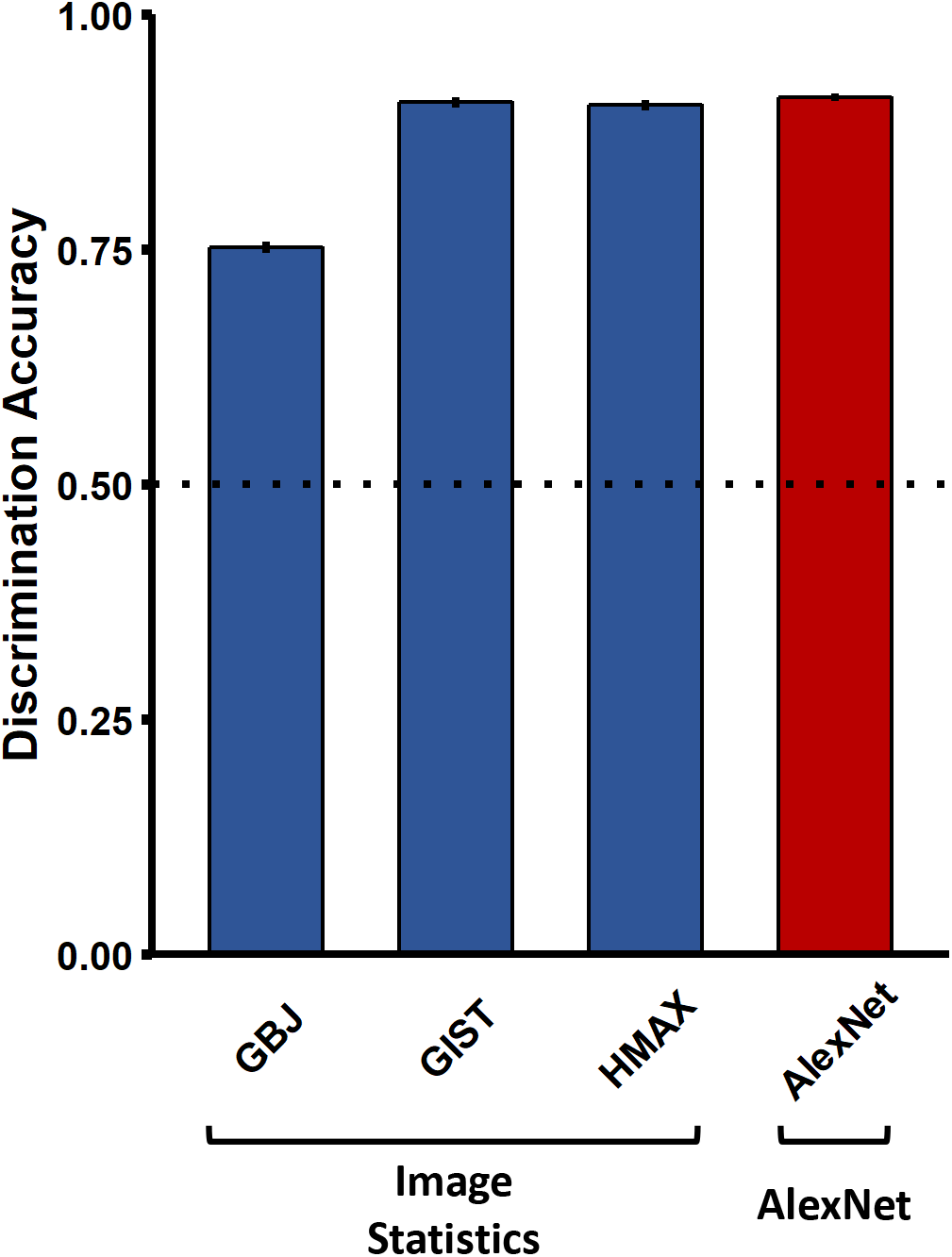
Discrimination accuracies for all non-skeletal models. Each model was able to discriminate between objects significantly above chance (dotted line).

**Supplemental Figure 2.**
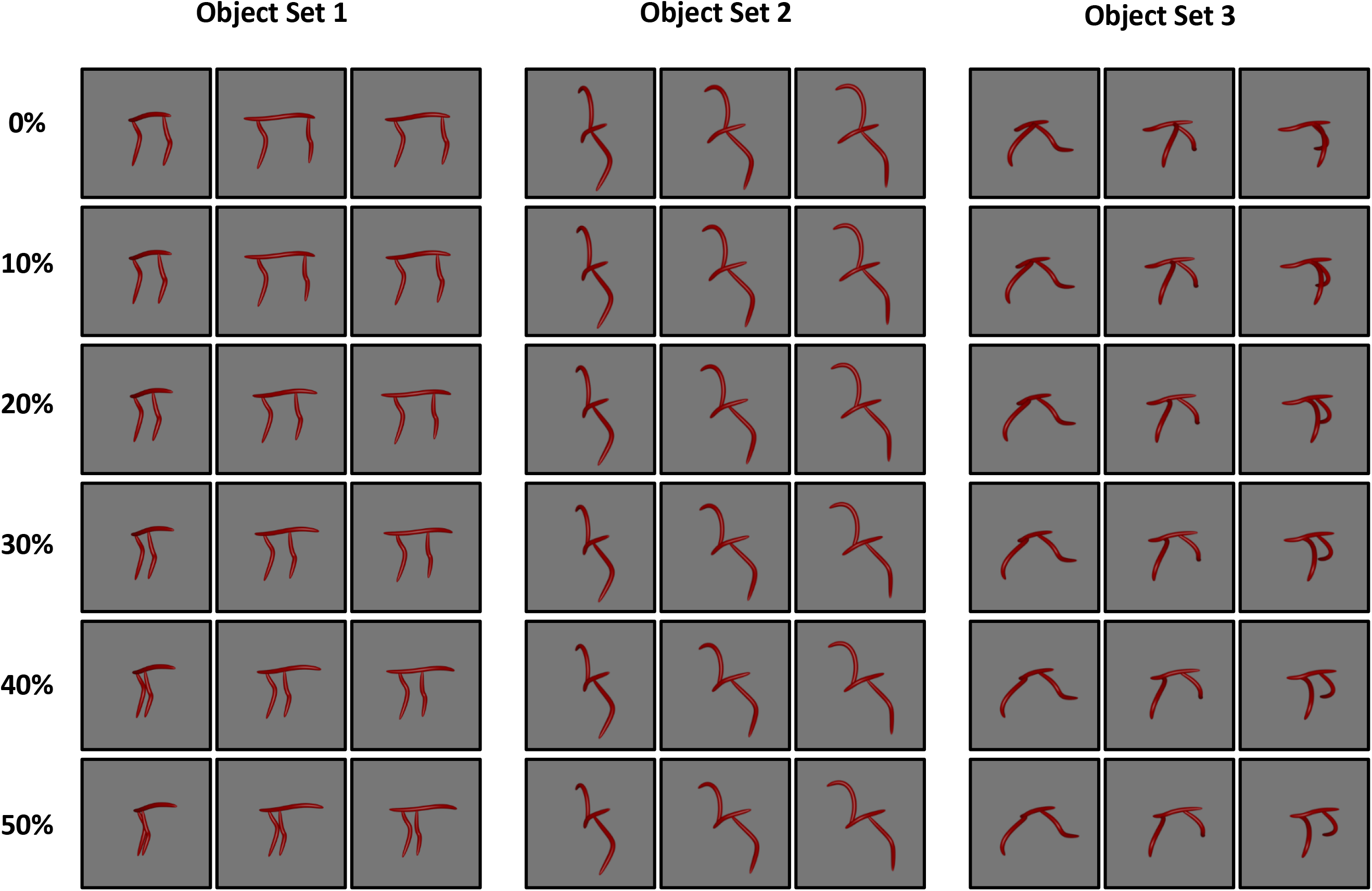
All stimuli used in Experiment 2. Objects were comprised of three sets, each with distinct coarse spatial relations. Within each set, objects varied in skeletal similarity by increments of 0%, 10%, 20%, 30%, 40%, or 50% (each row). Each object could be presented in one of three orientations, each of which is depicted here (30°, 60°, 90°; each column within an object set).

**Supplemental Figure 3.**
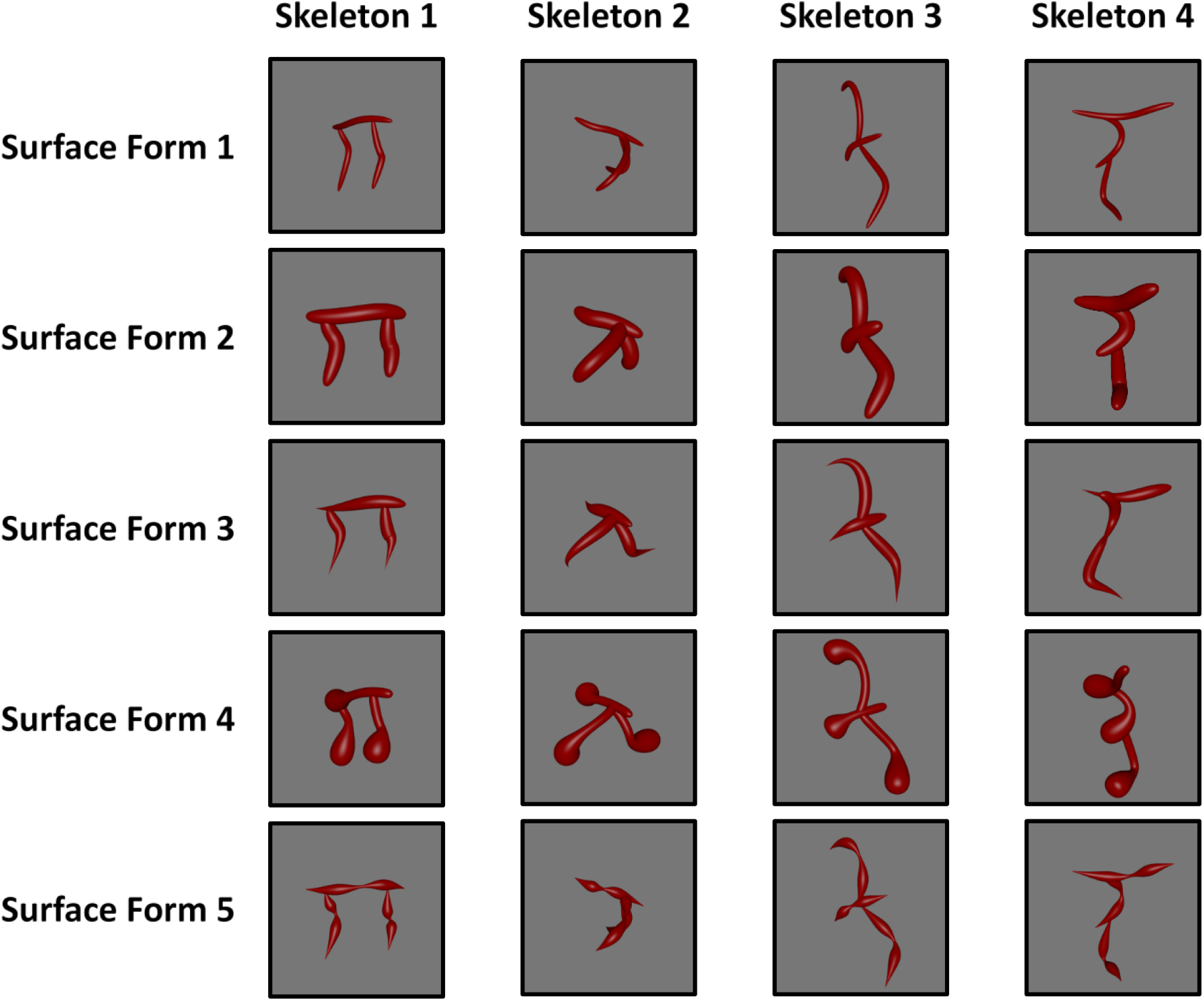
The stimulus set used in Experiment 3. Each column displays objects with the same skeleton, but different surface forms. Each row displays objects with the same surface form, but different skeletons. Each object could be presented in one of three orientations (30°, 60°, 90°), a subset are depicted here.

## Author Contributions

VA performed the experiments and analyzed the data. VA and SFL wrote the manuscript. Both authors conceived and designed the experiments, interpreted the experimental results, and approved the submission.

